# Nuclear myosin I mediates genomic clustering of estrogen receptor-α without affecting ligand-induced transcriptional robustness

**DOI:** 10.1101/2025.01.29.635522

**Authors:** Sudha Swaminathan, Ishfaq Ahamd Pandith, Rajat Mann, Deepanshu Soota, Arif Hussain Najar, Dimple Notani

## Abstract

Ligand-induced transcriptional responses are rapid and robust. Estrogen, one such ligand, exerts its rapid effects through the steroid hormone receptor estrogen receptor-alpha (ERα). Upon stimulation, ERα binds to clustered enhancers to drive target gene expression. Here, we identify nuclear myosin 1 (NM1/Myo1c) as a positive regulator of ERα clustering on enhancers, promoting condensate formation on chromatin. NM1 is enriched on chromatin at the peak of the signaling response, and its depletion leads to a genome-wide reduction in ERα occupancy and condensates. Surprisingly, these alterations in chromatin occupancy of ERα have minimal impact on estrogen-regulated gene expression, suggesting that transcriptional output remains robust despite disruptions in transcription factor clustering. This study reveals a regulatory axis between Myo1c and ERα clustering, offering fresh insights into the intricacies of transcriptional regulation.

## Introduction

The transcriptional response to signalling is inherently fast and dynamic, necessitating active processes to drive its inducible nature. Acute transcriptional induction hinges on the robust recruitment of transcriptional machinery and polymerase complexes to target regions in the genome. This is accompanied by the clustering of genomic elements and components of the transcriptional machinery. These extensive movements of chromatin, polymerases, and other proteins enable, both their assembly and disassembly at the start and end of signalling respectively.

Interestingly, nuclear myosins and other cytoskeletal proteins (like Rac and actin) associate with the transcriptional machinery in the nucleus specifically during signalling (Kysela et al., 2005; Majewski et al., 2018; Sun et al., 2021; Knerr et al., 2023). Remarkably, in diverse models of induction like differentiation, antigenic stimulation, or chemical stimulation, nuclear myosins interact with the transcription machinery (Zorca & Kim et al., 2015; Majewski et al., 2018; Xue et al., 2019). However, their exact functions remain poorly understood (de Lanerolle, 2012; Fili & Toseland, 2020). In this context, alongside polymerases, chromatin remodellers, and loop extrusion machinery, the nuclear pool of actin and myosins may perform critical motor functions by regulating the movement of chromatin, transcription factors, and transcriptional machinery to enable rapid transcriptional induction.

Myosin1c (Myo1c) is a cytoskeletal protein that performs motor and tether-like functions in the cytosol. It moves molecular cargo (e.g. vesicles) over actin filaments in cytoplasm and tethers adaptor proteins in the plasma membrane to the underlying actin network (Bond et al., 2013). Interestingly, it was the first myosin to be detected in the nucleus (Nowak et al., 1997; Pestic-Dragovich et al., 2000). Myo1c has three isoforms, and Nuclear Myosin I (NM1) is the isoform localizing in the nucleus. Since nuclear actin is very transient, dynamic, and challenging to track as opposed to cytosolic actin (Baarlink et al., 2013; Plessner et al., 2015), it is difficult to ascertain if myosins perform motor or tether-like functions in the nucleus. The steroid-responsive nuclear receptor, estrogen receptor alpha (ERα) interacts with Myo1c in the nucleus upon estrogen stimulation (Ambrosino et al., 2010). Multiple studies have confirmed this interaction by co-immunoprecipitation and mass spectrometry (Tarallo et al., 2011; Marsh et al., 2017). However, the biological function and significance of this interaction remain unknown.

ERα has dual functions: it is a receptor for estrogen hormone (17β-estradiol or ‘E2’), and also a transcription factor (TF) (O’Malley & Tsai, 1992). Upon binding to E2, ERα dimerizes, moves into the nucleus, and binds to specific estrogen response elements (EREs) in the DNA. ERα binding promotes recruitment of coactivators and RNA Polymerase II on these genomic loci, which ultimately leads to the transcription of these regions (Tsai & O’Malley, 1994; Green & Carroll, 2007). Recent studies show that liganded ERα binds and assembles enhancer clusters in 3D and undergoes phase-separation (Bojcsuk et al., 2017; Saravanan et al., 2020; Boija & Klein et al., 2018; Nair et al., 2019).

In this regard, the known interaction of ERα and Myo1c invokes a possibility of collaboration of TF and nuclear motors in E2-induced transcription. To test the role of this interaction, we depleted Myo1c in MCF-7 cells and observed a drastic loss of ERα binding genome-wide upon E2 stimulation. The loss of binding was recapitulated by a reduction in ERα condensates. Despite these binding alterations, the estrogen-induced gene transcription was unaltered, reflecting the robustness of the transcriptional program. These findings reveal the unappreciated role of NM1 in transcription factor binding and condensate formation.

## Results

### Genomic occupancy of ERα is dependent on Myo1c

In the cascade of E2 signalling, binding of ERα to EREs denotes an important initial step (O’Malley & Means, 1974; O’Malley & Tsai, 1992). Since Myo1c and ERα are reported to interact with each other, we wanted to investigate how Myo1c might regulate E2-induced ERα binding. We standardized siRNA-mediated knockdown of Myo1c in MCF-7 cells (see Methods) and validated the efficiency of knockdown (Figures 1A and S3A). ERα genomic occupancy is influenced by concentration of ERα protein, so we checked if the knockdown of Myo1c affected ERα levels. We found that ERα protein levels, in whole cell lysates as well as in nuclear fraction, were unaffected by Myo1c knockdown (Figures 1A and S1A). Subcellular fractionation also confirmed that Myo1c knockdown does not perturb the E2-induced translocation of ERα from cytosol to nucleus (Figure S1A). Thus, Myo1c knockdown does not change total ERα protein levels and its distribution in cytosol and nucleus.

**Figure 1.**
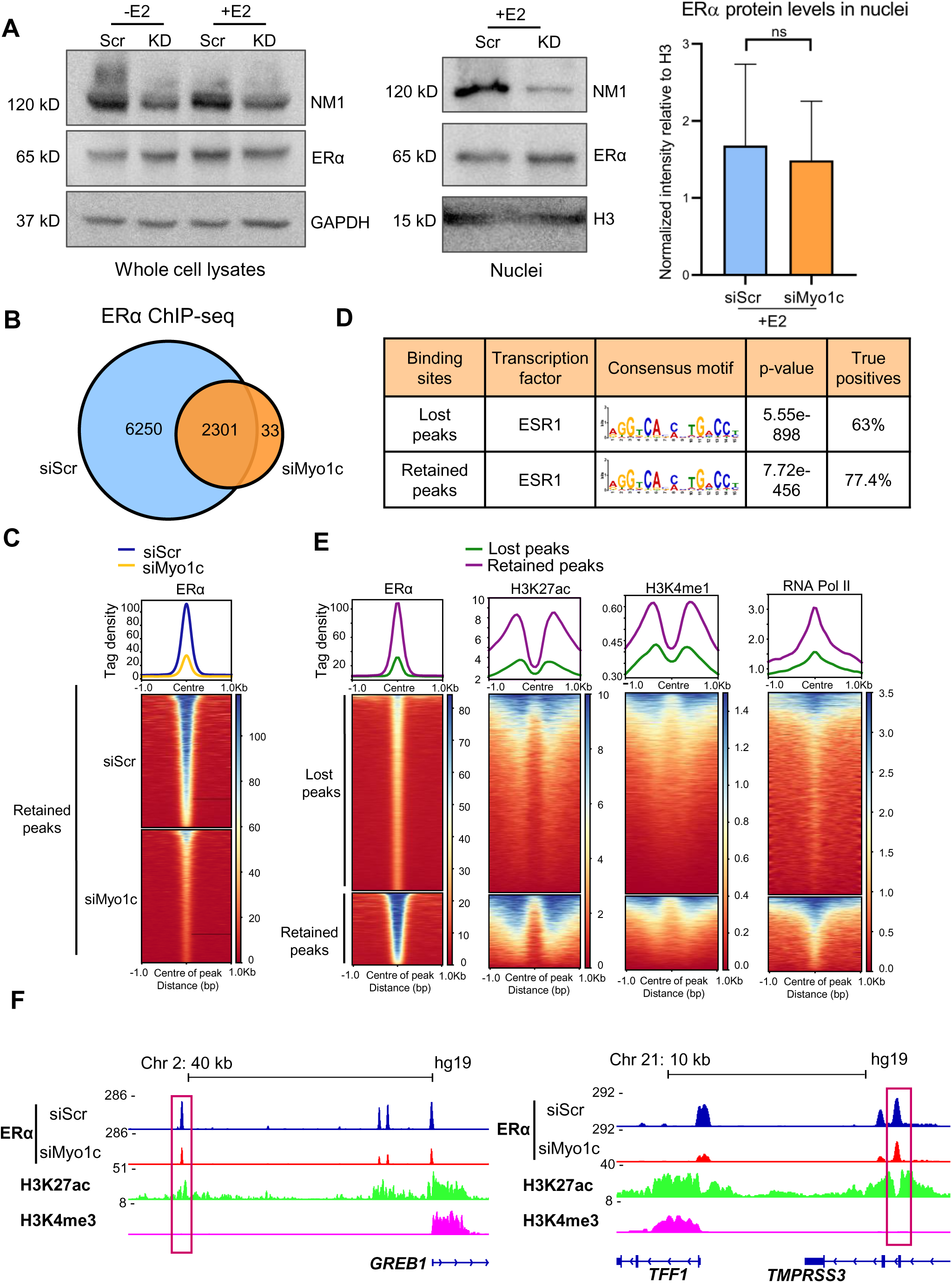
Genomic occupancy of ERα is dependent on Myo1c. (A) Western blots showing ERα protein levels in whole cell lysates and nuclear fractions of siScr (‘Scr’) and siMyo1c (‘KD’) samples (left and middle panels); - E2: vehicle treatment; +E2: E2 treatment. Bar graph showing nuclear ERα levels do not change upon Myo1c knockdown (right panel). Background-normalized band intensities of ERα relative to those of H3 histone are plotted. Paired t-test was used to test significance (p-value = 0.3926); whiskers indicate standard deviation around mean; N=3. (B) Venn diagram showing ERα peaks detected in ERα ChIP-seq in siScr and siMyo1c samples at 60 minutes of E2 stimulation. (C) Heatmap showing ERα ChIP-seq signal decreases even at the retained peaks in the siMyo1c sample, as compared to siScr sample. (D) Table depicting motifs with highest enrichment score in the different categories of peaks shown in panel B. (E) Heatmaps showing ChIP-seq signals for ERα, H3K27ac, H3K4me1 and RNA Pol II upon E2 treatment are higher at the ‘retained’ ERα peaks, as compared to the ‘lost’ ERα peaks. ERα signal intensity was plotted using siScr sample. Heatmaps for H3K27ac, H3K4me1 and RNA Pol II were generated from published datasets (see Methods). (F) IGV screenshots showing loss of ERα ChIP-seq signal upon Myo1c knockdown at enhancer clusters regulating *GREB1* gene and *TFF1* gene. Major enhancers are highlighted in red boxes. All tracks show ChIP signal from MCF-7 cells post 60 minutes of E2 stimulation; H3K27ac and H3K4me3 tracks were generated using published datasets (see Methods).

In order to test if Myo1c regulates ERα occupancy on chromatin, we performed ERα ChIP-seq in control (si-scrambled; ‘siScr’) and Myo1c-knockdown cells stimulated by E2 for 1 hour in two biological replicates. We detected a total of 8584 ERα peaks (Figures 1B and S1B; see Methods for details on peak calling). In both replicates, we observed a profound decrease in the number of ERα peaks upon Myo1c knockdown (Figures 1B and S1B). About 2300 peaks still remained after Myo1c knockdown but they exhibited very low mean normalized read counts compared to the control sample (Figures 1C and S1C). This indicates an overall decrease in ERα binding at all sites upon Myo1c knockdown. All peaks were bonafide ERα binding sites as the estrogen receptor motif was the top enriched motif in them (Figures 1D and S1E). Further, ‘retained’ peaks were inherently stronger binding sites for ERα, as they showed higher ERα binding in the control samples than the ‘lost’ sites (Figures 1E and S1D). These ‘retained’ sites also exhibited higher levels of H3K27ac, H3K4me1 and RNA Pol II (Figure 1E). The data suggests that Myo1c positively regulates ERα binding in the genome.

ERα-regulated transcriptional response is driven by ERα-bound enhancers. These enhancers show clustered binding of ERα upon estrogen stimulation (Saravanan et al., 2020). A closer view at these enhancers at classical E2-responsive genes, like *TFF1* and *GREB1*, reveals the acute loss of ERα binding upon Myo1c depletion (Figures 1F and S1F). Thus, even though levels of ERα protein in the nucleus do not change upon Myo1c knockdown, there is a dramatic reduction in its binding genome-wide.

### Myo1c depletion perturbs ligand-dependent ERα condensates

ERα forms condensates in the nucleus post E2 stimulation (Boija & Klein et al., 2018; Nair et al., 2019; Saravanan et al., 2020). These condensates are formed on ERα-bound clustered enhancers which regulate genes such as *TFF1* (Saravanan et al., 2020). Since Myo1c depletion affected ERα binding on regulatory elements, we investigated if Myo1c knockdown also affects the formation of ERα condensates using immunofluorescence (Figure 2). We treated the control and Myo1c-knockdown cells with E2 (or ethanol; see Methods for details) for 1 hour. Then, we treated them with cytoskeletal buffer to remove nucleoplasm-soluble (or non-chromatin-bound) proteins before fixation (Fey et al., 1984; Jackson et al., 1988; Fujita et al., 1997; Okuno et al., 2001; Groth et al., 2007; Turgay et al., 2014). This was done to ensure that only chromatin-bound ERα is detected by immunofluorescence.

**Figure 2:**
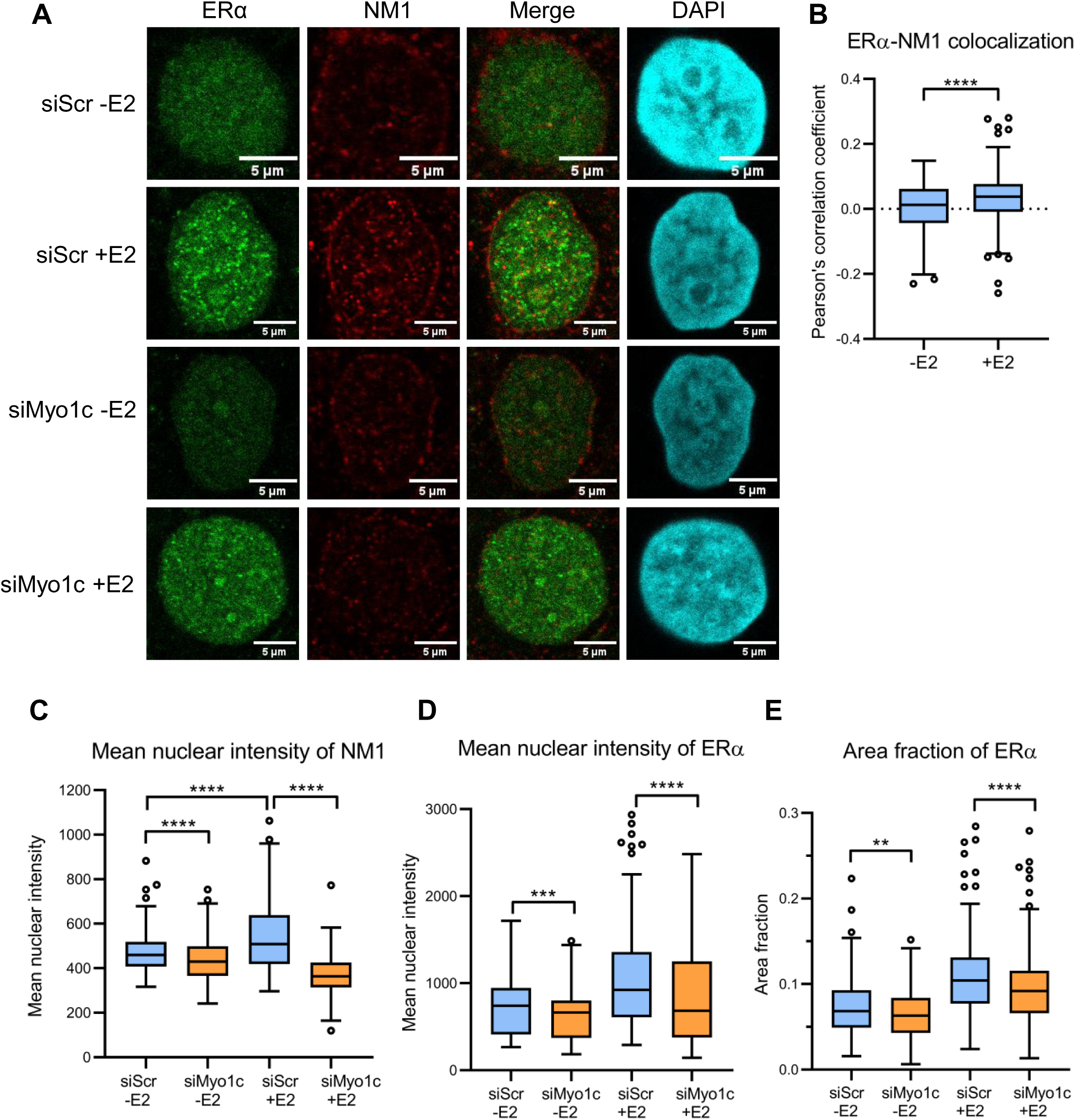
Myo1c depletion perturbs ligand-dependent ERα condensates. (A) Confocal images of immunofluorescence for ERα (green) and NM1 (red) in CSK-treated samples are shown. Chromatin-bound ERα and NM1 foci are most evident in siScr +E2 condition. DAPI is shown in cyan. Scale bar: 5 μm. -E2: vehicle treatment; +E2: E2 treatment for 60 minutes. (B) Pearson’s correlation coefficient between ERα and NM1 signals increases upon E2 treatment. Significance was tested using Mann-Whitney test. Centre lines in boxes indicate medians, whiskers extend to 1.5 times the IQR from 25^th^ and 75^th^ percentiles (Tukey), **** indicates p-value<0.0001. N=3. Number of nuclei per sample: 298, 281 respectively. (C) Validation of Myo1c knockdown by quantification of NM1 mean nuclear intensity. Mann-Whitney test was performed to test significance. Centre lines in boxes indicate medians; whiskers extend to 1.5 times the IQR from 25^th^ and 75^th^ percentiles (Tukey); **** indicates p-value<0.0001; N=3. Number of nuclei per sample: 298, 273, 281, 278 respectively. (D) Chromatin-associated ERα foci decrease upon Myo1c knockdown. Mean nuclear intensity of chromatin-bound ERα and (E) area fraction of chromatin-bound ERα foci decrease upon Myo1c knockdown. For panels D and E: Mann-Whitney test was performed to test significance. Centre lines in boxes indicate medians; whiskers extend to 1.5 times the IQR from 25^th^ and 75^th^ percentiles (Tukey); **** indicates p-value<0.0001; *** indicates p-value <0.001; ** indicates p-value<0.01. N=3. Number of nuclei per sample: 298, 273, 281, 278 respectively.

ERα translocates to the nucleus and binds to chromatin upon E2 stimulation. Thus, its signal was higher in the E2-treated cells than vehicle-treated control cells (Figure 2A). Unexpectedly, we found that E2 stimulation leads to an increase in NM1 protein in the nucleus within 60 minutes (Figure 2A and S2). There are examples of cytoskeletal proteins translocating to the nucleus when cells are stimulated by hormones/chemicals (Samstag et al., 1994; Majewski et al., 2018). A recent study shows that serum stimulation in HeLa cells not only leads to appearance of myosin VI foci in the nucleus, but also an increase in colocalization of myosin VI foci with RNA Pol II, indicating the interaction of nuclear myosin VI with transcription machinery (Hari-Gupta et al., 2022). In line with this, we found that colocalization of chromatin-bound ERα and NM1 was higher in E2-treated samples (Figure 2B). Since levels of both proteins were increasing in the nucleus upon E2 stimulation, we tested if association of ERα to chromatin is dependent on Myo1c.

We calculated the mean nuclear intensity of NM1 in all samples, and as per expectation, NM1 was significantly less in the knockdown samples than the control samples (Figure 2C). Next, we tested if similar was also true for ERα between control and Myo1c-knockdown samples. Indeed, we found that the mean nuclear intensity of chromatin-bound ERα was also lower in the Myo1c-knockdown cells (Figure 2D). This suggests that Myo1c knockdown leads to reduction in the amount of ERα bound to chromatin, which was reflected in ERα ChIP-seq (Figure 1 and S1). In addition, ERα forms condensates on E2-regulated genes (Saravanan et al., 2020). In corroboration with ChIP-seq which shows loss of ERα on clustered enhancers (Figure 1F and S1F), the area fraction of chromatin-bound ERα foci was lower in the Myo1c-knockdown cells (Figure 2F). This suggests that chromatin-bound ERα forms smaller foci in the background of Myo1c knockdown as compared to control.

Overall, the microscopy data suggests that Myo1c associates with chromatin at the peak of signalling where it co-localises with ERα condensates. Consolidating these observations with results of ChIP-seq, we infer that that Myo1c is important for the association of ERα with chromatin and the resultant condensates upon E2 stimulation.

### Transcription of E2-induced genes is unaffected by Myo1c knockdown

We proceeded to check how loss of ERα binding on chromatin might impinge on the transcription of E2-target genes. We performed RNA-seq in control and Myo1c-knockdown cells under vehicle-treated and E2-treated conditions. We first verified E2-mediated activation of gene transcription comparing the vehicle- and E2-treated samples (Figure 3A). Consistent with the results of western blotting (Figure 1A), we found that Myo1c knockdown did not change the levels of estrogen receptor (*ESR1*) transcripts (Figure S3B). In order to test gene transcription perturbations upon Myo1c knockdown, we compared siScr and siMyo1c conditions. Surprisingly, we did not detect any changes in the expression of total (mature RNA) as well as intron-mapped transcripts (unspliced RNA) of known E2-induced genes upon Myo1c-knockdown (Figures 3B-D, S3 C-D; see ‘Methods’ for details).

**Figure 3:**
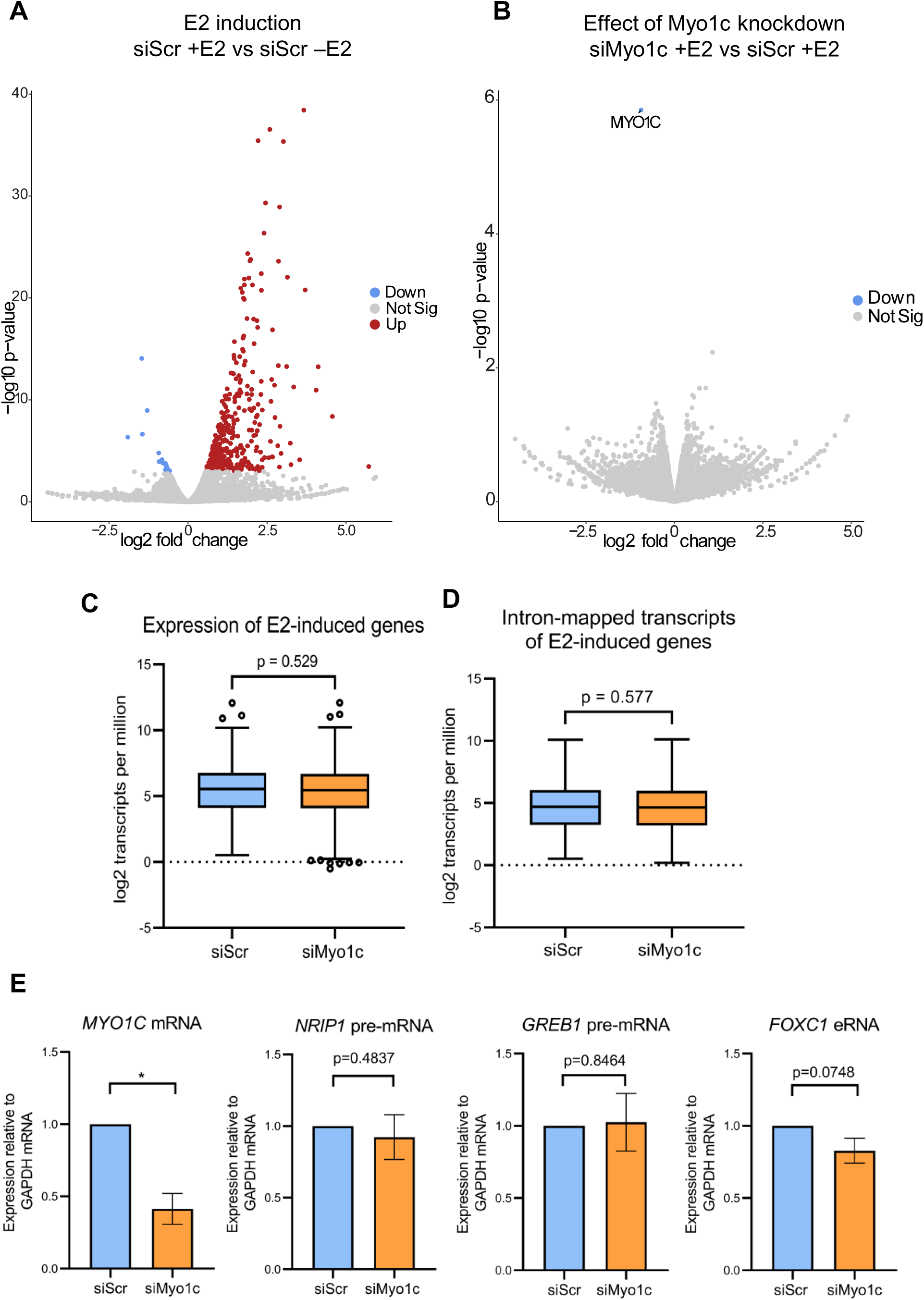
Transcription of E2-induced genes is unaffected by Myo1c knockdown. (A) Confirmation of E2 induction. Volcano plot shows genes with significant p-value (red and blue dots) in siScr +E2 over siScr -E2 samples. Values were obtained from DESeq2 using two biological replicates of RNA-seq. (B) Volcano plot showing knockdown of Myo1c in siMyo1c +E2 sample relative to siScr +E2 sample. Values were obtained from DESeq2 using two biological replicates of RNA-seq. (C) Steady-state transcript levels of 1200 top E2-induced genes (Hah et al, 2011) from total RNA-seq. (D) Quantification of transcripts of E2-induced genes (same as panel C) mapping to introns. P-values in (C) and (D) were calculated using Mann-Whitney test. Centre lines within boxes represent medians. Whiskers represent values within 1.5 IQR (Tukey). (E) Expression of Myo1c mRNA and nascent RNA of classical E2-responsive genes at 1 hour of E2 stimulation quantified by RT-qPCR. P-values were calculated by paired t-test; * represents p-value < 0.05; whiskers represent standard deviation around mean; N=3.

**Figure 4:**
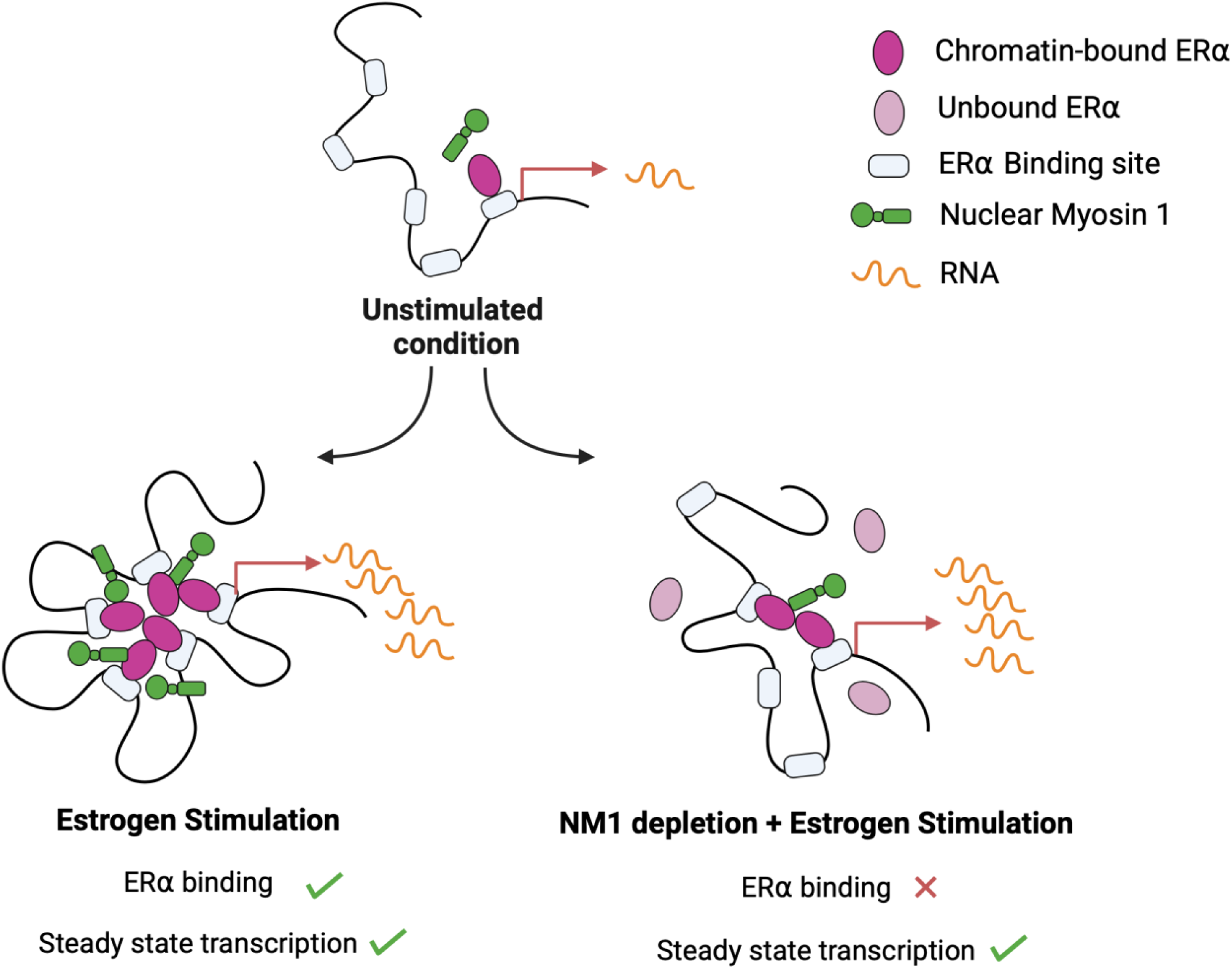
Model showing regulation of ERα binding in the genome by NM1. In unstimulated condition, very few EREs in the genome are bound by ERα. Upon E2 stimulation, liganded ERα molecules bind to EREs genome-wide. NM1 levels in the nucleus increase and NM1 molecules colocalize with ERα condensates. ERα recruits transcription machinery and target genes are transcribed. However, depletion of NM1 impairs this signalling-induced association of ERα with chromatin. Though chromatin-bound ERα forms smaller condensates, steady state transcription remains stable. Schematic created using BioRender.

To confirm the unexpected results of RNA-seq, we used qRT-PCRs to test the levels of nascent transcripts of classical E2-induced genes. We found that Myo1c knockdown, albeit decreasing the amount of chromatin-bound ERα, has minimal to no effects on transcription of E2-target genes (Figure 3E). In corroboration with this, recent studies also show that low levels or reduced chromatin binding or smaller condensates of ERα do not lead to reduced transcription (Bohra & Islam et al., 2024; Soota et al., 2024; Henninger & Oksuz et al., 2021). A very recent study points to how differences like loss of co-regulation of E2-induced genes in single cells are not captured by routinely-used techniques like RNA-seq (Dong et al., 2024). Population-level assays like qRT-PCRs and RNA-seq may buffer the differences present at the single cell level, thereby masking the underlying biology (Dong et al., 2024). Thus, we deduced that changes in chromatin-association of ERα may not either directly translate into changes in transcription at 1 hour of E2 stimulation or the modest changes are masked by bulk amplification/sequencing.

## Discussion

Cytoskeletal proteins are part of transcription complexes that assemble on chromatin in response to specific ligand stimulation (Kysela et al., 2005; Zorca & Kim et al., 2015; Xue et al., 2019; Sun et al., 2021). NM1 is a known interacting partner of nuclear ERα (Ambrosino et al., 2010; Tarallo et al., 2011; Marsh et al., 2017). In this study, we have delineated the role of nuclear myosin 1 (NM1/Myo1c) in E2-induced signalling. Using siRNA-mediated knockdown approach, we observe that NM1 regulates ERα binding in the genome. Knockdown of NM1 results in genome-wide reduction in ERα occupancy (Figure 1), indicating that the presence of NM1 protein assists in binding of ERα to chromatin. This finding is supported by microscopy data which show decrease in chromatin-bound ERα condensates upon NM1 knockdown (Figure 2). However, the loss of condensates does not affect estrogen-induced steady state transcription.

The relationship between ERα binding on DNA and its functional consequences on transcription is complex, particularly when influenced by factors like the motor protein myosin. While it is unclear how myosin might directly affect ERα-DNA interactions, intriguing observations emerge when examining transcriptional outcomes. Upon knockdown of NM1, a protein implicated in inducible transcription (Ambrosino et al., 2010; Kysela et al., 2005), a dramatic loss of ERα binding is observed, as evident from ChIP-seq and microscopy data. Despite reduction in transcription factor (TF) binding, ligand-dependent gene transcription (nascent RNA) appears unaffected. This paradox suggests that the remaining TF binding, even if reduced, might be sufficient to sustain transcription. Other studies also illustrate that transcription factor binding and transcript levels in the population may not be well-correlated (Soota et al., 2024; Henninger & Oksuz et al., 2021). In a given population, cells exhibit a range of ERα levels, and therefore vary in the amount of ERα protein bound to chromatin. Very recent work by Bohra & Islam et al. (2024) presents evidence for the non-intuitive relationship between ERα levels and nascent transcription in single cells. The authors show that even cells with low levels of ERα produce as many nascent transcripts of *TFF1* gene as cells containing median levels of ERα at 1 hour post E2 stimulation (Bohra & Islam et al., 2024). This nuanced relationship highlights the resilience of transcriptional systems and suggests that optimal transcription (nascent RNA) is maintained even in the face of molecular perturbations like NM1 knockdown.

Further, another myosin present in nucleus, myosin VI also interacts with nuclear receptors, however, its knockdown does not lead to a general decrease in E2-regulated gene expression (Fili et al., 2017). Similarly, nuclear myosin VI depletion in a different inducible system of T-cells perturbs transcription elongation without affecting the bulk levels of mature mRNA produced from the induced genes (Zorca & Kim et al., 2015). In fact, very recently, two independent studies investigating the effects of depletion of nuclear myosin and nuclear actin found dramatic changes in genome architecture and accessibility respectively, but observed very little correlated changes in transcription by RNA-seq (Peng et al., 2024; Sen et al., 2024). Thus, overall transcriptional output (mature mRNA) may remain relatively stable through population-level dynamics despite a drastic reduction in nuclear myosin levels across the population.

Importantly, bulk assays like RNA-seq and qRT-PCR fail to capture differences occurring at the single-cell level (Dong et al., 2024). The study by Dong et al. (2024) elucidates how changes in genome architecture by loss of cohesin could change the pairs of genes that are co-expressed in a cell but these changes are not evident at the population level. Connecting our observations with these findings, it is possible that changes in NM1 levels could alter gene expression at the single cell level. Our observations in qRT-PCRs of nascent transcripts and bulk RNA-seq could be explained by a) low levels of chromatin-bound ERα being sufficient to sustain nascent transcription or b) masking of single cell heterogeneity due to usage of population-level measurements of mature transcripts.

Our study adds to the growing field of ligand-induced gene regulatory control by cytoskeletal proteins in the nucleus (Zorca & Kim et al., 2015; Sun et al., 2021; Knerr et al., 2023). Our findings point to how presence of nuclear myosin is critical for optimal ERα binding in vivo. To our knowledge, this is the first report of transcription factor binding to chromatin being modulated by nuclear myosin. This modulation may occur at the level of individual binding events of ERα on EREs, or change the duration for which ERα condensates occupy EREs, or it may enhance the clustering of ERα-bound sites. More experiments are needed to unravel the underlying mechanism. Single cell sequencing and super-resolution imaging may provide answers to how cytoskeletal proteins in the nucleus regulate transcription factor dynamics and nascent transcription.

## Materials & Methods

### Cell culture

MCF-7 human breast adenocarcinoma cells were cultured in complete media, i.e., Dulbecco’s Modified Eagle Medium (10569-010, Gibco) supplemented with 10% FBS (10437028, Gibco) and 1% penicillin-streptomycin (‘pen-strep’; 15140122, Gibco) at 37 degrees Celsius and 5% CO2 in humidified incubator. Cells were split using 1X Trypsin-EDTA at 72-96 hours post seeding and reseeded at 1/4th confluence for maintenance.

### siRNA transfections and E2 treatment

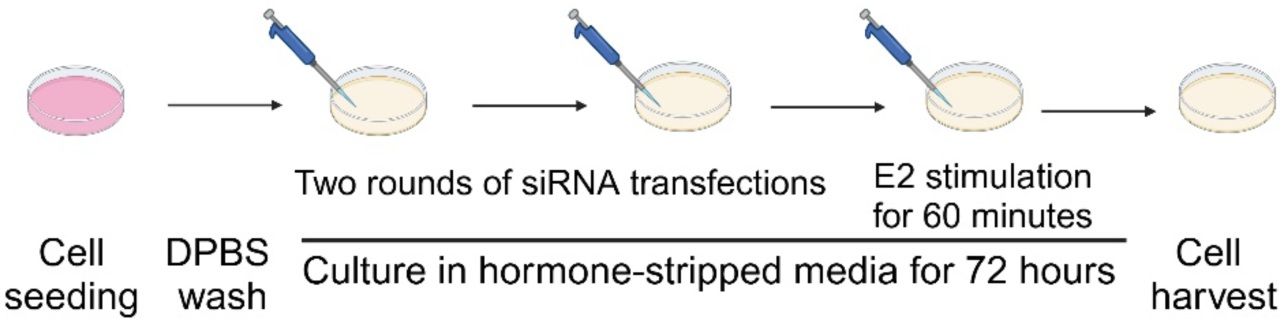

ON-TARGETplus (Dharmacon Horizon Discovery) siRNAs (Non-targeting Control Pool, D-001810-10-20 and against human Myo1c, L-015121-00-0005 SMARTPool) were resuspended in Ambion nuclease-free water (AM9937, Invitrogen) as recommended in Horizon Discovery manual to obtain 20 micromolar stock and stored at −20 degree Celsius until use. siRNA transfections were performed twice, with the final concentration of siRNA added to each 10 cm plate in each round being 10nM. For siRNA transfections, cells were seeded at 30% confluence in 10 cm dishes in complete media. After 16 hours of seeding, they were washed once with 1X DPBS (no calcium, no magnesium;14190-144, Gibco) and transfected with 2.5 µl siRNA dissolved in 1 ml reduced serum OptiMEM without phenol red (11058-021, Gibco). 15µl Lipofectamine 3000 (L3000-008, Invitrogen) was used per 10 cm dish. Final volume of OptiMEM in each 10 cm dish was 5ml. Media was replaced by 10 ml stripping media, i.e., DMEM (no FBS, no phenol red; 21063-029, Gibco) supplemented with 5% charcoal-stripped FBS (A33821-01, Gibco) and 1% pen-strep (15140122, Gibco) after 6 hours. Second transfection was performed in the same way at 24 hours post the first transfection. Media was replaced by 10 ml stripping media after 6 hours. Cells were subjected to vehicle (crude ethanol) or 100 nM beta estradiol (‘E2’; E2758, Sigma Aldrich) after 72 hours of culture in hormone-deprived media. After 60 minutes of treatment, cells were harvested for RT-qPCR, SDS-PAGE, or ChIP.

For immunostaining, transfection mix volumes were used as per recommendation for 6-well plates. Volumes of siRNA and Lipofectamine 3000 were 0.5µl and 6 µl per well respectively. Final volume of OptiMEM (without phenol red) per well was 1 ml. The final concentration of siRNA added to each well in each round was 10nM. After 72 hours of culture in hormone-deprived media, cells were stimulated with ethanol or 100 nM E2 for 60 minutes.

### Chromatin immunoprecipitation (ChIP)

Cells cultured in 10 cm plates (∼20 million cells) were crosslinked in 1% formaldehyde for 10 minutes at room temperature and then treated with 0.125M glycine for 5 minutes. Cells were washed thrice with ice-cold 1X PBS to remove traces of formaldehyde and glycine. They were scraped and pelleted in 1X ice cold PBS by centrifugation at 3000 rpm at 4 degrees Celsius for 6 minutes. Cell pellets were stored at −80 degrees Celsius overnight. Cell pellets were thawed on ice and lysed with L2 Nuclear Lysis Buffer (50 mM Tris-HCl pH 7.4, 1% SDS, 10 mM EDTA) containing 1X Protease Inhibitor Cocktail. After incubation on ice for 15 minutes, lysates were sonicated for 28 cycles (30s ON and 30s OFF per cycle) using Bioruptor Pico (Diagenode) to obtain fragments in the range of 400 to 800 bps. Sonicated lysates were centrifuged at 12000 rpm at 4 degrees Celsius for 12 minutes. Supernatant was collected and concentration of DNA was measured. Supernatant was diluted 2.5 times using dilution buffer (20 mM Tris-HCl pH 7.4, 100 mM NaCl, 2 mM EDTA pH 8.0, 0.5% Triton-X-100). 100 µg chromatin was used for each IP and beads sample. Input samples were set up with 1% of chromatin (10 µg). 1µg ERα (sc-8002, Santa Cruz) mouse monoclonal antibody was added to each IP sample. Tubes were incubated at 4 degrees Celsius overnight at low-speed rotation. Protein G Dynabeads (10004D, Invitrogen) were washed thrice with 1X PBS, blocked with 5% BSA and washed thrice with 1X PBS. 15 µl of the beads was added to each of the beads and IP samples. Samples were incubated at 4 degrees Celsius at low-speed rotation for 4 hours. Complexes were captured on magnetic rack and washed sequentially with wash buffers I, II, III and 1X Tris-EDTA for 5 minutes at 4 degrees at high-speed rotation. Compositions of wash buffers were as follows: Wash buffer I: 20 mM Tris-HCl pH 7.4, 150 mM NaCl, 0.1% SDS, 2 mM EDTA pH 8.0, 1% Triton-X-100; Wash buffer II: 20 mM Tris-HCl pH 7.4, 500 mM NaCl, 2 mM EDTA pH 8.0, 1% Triton-X-100; Wash buffer III: 10 mM Tris-HCl pH 7.4, 250 mM LiCl, 1% NP-40, 1% sodium deoxycholate, 1mM EDTA pH 8.0. Complexes were eluted in fresh elution buffer (100 mM sodium bicarbonate, 1% SDS) at 37 degrees Celsius for 30 minutes at 1400 rpm in thermomixer. Reverse crosslinking was performed in 5M NaCl overnight at 65 degrees Celsius and 600 rpm in thermomixer. Solutions were subjected to phenol-chloroform-isoamyl alcohol-based method to purify DNA. Purified DNA was dissolved in 10 µl nuclease-free water per input, beads and IP sample and stored at −20 degrees Celsius.

### ChIP-sequencing and analysis

*For ERα ChIP-seq in siScr and siMyo1c samples:* ChIP-seq libraries were prepared using NEBNext® Ultra™ II DNA Library Prep with Sample Purification Beads (E7103L) as per manufacturer’s guidelines. The amplified libraries were size-selected to enrich for fragments in the range of 300 to 500 bp. Single-end sequencing was performed in 1×50 bp format on HiSeq 2500 (Illumina Inc.). Sequenced reads were uploaded to Galaxy web platform and analyzed using the resources available on the public server at usegalaxy.org or usegalaxy.eu (Jalili et al., 2020). Firstly, quality of sequencing reads was affirmed using FastQC. Unmapped reads and reads with low Phred score were removed using FASTQ Groomer. Then, reads were aligned to human genome hg19 version (GRCh37) using Bowtie2 with default settings. Unmapped reads, reads with low mapping score (<20) and duplicate reads were removed by filtering the BAM files. In one of the replicates, 1% spike-in of *Drosophila* genomic DNA was added at the time of sequencing. Reads mapping to *Drosophila* genome were removed during filtering. Number of filtered human genome-mapped reads were almost equal between siScr and siMyo1c samples within each replicate. The filtered BAM files were processed with MACS2 *predictd* to determine average fragment size for peak calling (bandwidth for picking regions to compute fragment size was set as 300 bp; mfold range was set between 5 and 100). In case of replicates, *plotCorrelation* was used on multibamSummary files to ensure consistency among replicates. *plotFingerprint* was used to ensure that read count distribution among replicates was consistent. Narrow peaks were called with MACS2 *callpeak* after extending fragments to the sizes obtained from *predictd*. Peak calling was done without constructing a model (-- nomodel) and with a q value (FDR) set at 0.05. Input control file for MCF-7 cell line (E2-stimulated) was used from a published dataset with SRA accession number SRR350879 (Lee et al., 2012). This file was similarly processed with FASTQ Groomer and Bowtie2 to generate hg19-mapped BAM file, which was filtered to exclude unmapped, low-quality mapped and duplicate reads. Then, broad peaks were called for the input control dataset with MACS2 *callpeak* after extending fragments to the sizes obtained from MACS2 *predictd*. Peak calling was done without constructing a model (--nomodel) and with a q value (FDR) set at 0.05. hg19 blacklisted regions were removed from all peak files (ERα ChIP as well as input). Then, *bedtools SubtractBed* was used to subtract the filtered peaks obtained in input file from the filtered peaks obtained in each of the ERα ChIP-seq files. These were used as the final set of ERα peaks for all further analyses. Irreproducible discovery rate (IDR) analysis showed high consistency between peak ranks between siScr samples of both replicates. This consistency was lost when comparing peak ranks between siMyo1c samples of both replicates, further confirming the genome-wide loss of ERα binding occurring upon Myo1c knockdown. *Bedtools Intersect and SubtractBed* were used to identify peaks ‘lost’ and ‘retained’ in siMyo1c samples with respect to siScr samples. Bedgraph treatment files were converted into bigWig files for visualization in IGV browser. Datasets used for visualizing H3K27ac and H3K4me3 ChIP-seq signals as tracks in IGV were GSM2157740 and GSM3414774 respectively. For heatmap generation, bigWig files were used as score files in *computeMatrix* with the subtracted bed files of peaks as regions of interest. Matrix was constructed with the centre of each region set as reference point. Mean signal across each region (centred at midpoint) was plotted as heatmap with regions sorted in descending order using *plotHeatmap* tool.

#### Motif analysis

Motif enrichment was calculated using AME (MEME suite; McLeay & Bailey, 2010). ERα peaks called by MACS2 were filtered for blacklisted and input-enriched regions, as mentioned above. 100 bps flanking the summits of ERα peaks were used as primary sequences in AME. The corresponding shuffled sequences were used as background (control). HOCOMOCO Human (v11 core) database was used for motif identification. Average odds scores were calculated for the primary sequences and enrichment of motifs was tested through Fisher’s exact test in AME.

#### For H3K27ac, H3K4me1 and RNA Pol II ChIP-seq analysis from published datasets

To understand the distribution of H3K27ac, H3K4me1 and RNA Pol II on different sets of ERα peaks, we used following bigWig files from published datasets of E2-stimulated MCF-7 cells: GSM2157740 (H3K27ac), GSM3414780 (H3K4me1) and GSM2157760 (RNA Pol II). ERα peaks obtained after peak calling with MACS2 and filtering were used as regions of interest. *computeMatrix* was used to build a matrix with the centre of each region set as reference point. Mean signal across each region (centred at midpoint) was plotted using *plotHeatmap* with regions sorted in descending order.

### Subcellular fractionation

Approximately 10 million cells per sample were harvested on ice in 1X PBS. 100 µl of this suspension was mixed with 20 µl 6X Laemmli dye, boiled for 10 minutes to get whole cell lysates and stored at −20 degrees Celsius until use in SDS-PAGE. Rest of the cell suspension was centrifuged at 2500 rpm at 4 degrees to obtain cell pellets. Cell pellets were resuspended in Triton Extraction Buffer (1X PBS containing 0.5% Triton-X-100 and 1X Protease Inhibitor Cocktail) and incubated on ice for 15 minutes with gentle stirring. To obtain the cytoplasmic fraction, supernatant was collected after centrifugation at 3000 rpm for 5 minutes at 4 degrees Celsius. Cytoplasmic fraction was mixed with 6X Laemmli dye, boiled for 10 minutes and stored at −20 degrees Celsius until use in SDS-PAGE. The resultant nuclear pellets were washed thrice in 1X PBS containing 0.5% Triton-X-100 at 3000 rpm for 5 minutes at 4 degrees Celsius. Nuclear pellets were resuspended in 1X RIPA nuclear lysis buffer (50 mM Tris-HCl pH 7.4, 150 mM NaCl, 1% NP-40, 0.5% sodium deoxycholate, 0.1% SDS) and incubated on ice for 30 minutes with occasional mixing. The nuclear lysates were sonicated for 10 cycles (30s ON and 30s OFF per cycle) in Bioruptor Pico (Diagenode) at 4 degrees, mixed with 6X Laemmli dye, boiled for 10 minutes and stored at −20 degrees Celsius until use in SDS-PAGE.

### SDS-PAGE and western blotting

Samples were run on polyacrylamide gels (made from Surecast 40% acrylamide solution with 29:1 acrylamide:bis-acrylamide, Invitrogen; cast and run using Mini-PROTEAN tetra cell, Bio-Rad) and blotted onto 0.2 µm PVDF membranes (1620177, Bio-Rad) in transfer buffer (25 mM Tris, 0.192 M glycine and 20% methanol) using electrical transfer on ice (B1000, Mini Blot module, Invitrogen). Blots were probed with anti-GAPDH (mouse monoclonal antibody sc-32233, Santa Cruz Biotechnology), anti-ERα (mouse monoclonal antibody sc-8002, Santa Cruz Biotechnology) and anti-Myosin Iβ (Nuclear) rabbit antibody (M3567-200UL, Sigma) or anti-Histone H3 (rabbit antibody H0164, Sigma) antibodies in 5% non-fat dry milk in 1X TBST (tris-buffered saline with 0.1% Tween-20) overnight at 4 degrees Celsius. Blots were then washed thrice with 1X TBST for 5 minutes at room temperature at high speed. HRP-conjugated anti-mouse (1721011, Bio-Rad) or anti-rabbit (1706515, Bio-Rad) antibodies were used at 1:10,000 dilution in 1X TBST. Blots were incubated in the secondary antibody solution for 1 hour at room temperature and then washed thrice with 1X TBST for 5 minutes at room temperature at high speed. They were developed with Amersham enhanced chemiluminescence kit (RPN2106, Cytiva) and imaged with ImageQuant LAS 4000 GE. Images were analyzed with Fiji software (Schindelin et al., 2012). Band intensities were first background normalized, and then normalized to the appropriate loading control (GAPDH for total lysates as well as cytoplasmic fractions and, H3 for nuclear lysates respectively) in the same lane. Paired t-test was performed on data from three biological replicates to test significance.

### Immunostaining and image analysis

∼0.4 million cells per well were seeded in complete media in 6-well plates containing coverslips. After 16 hours, cells were washed with 1X DPBS and siRNA transfections were performed as mentioned above. After 72 hours of culture in hormone-deprived media, cells were stimulated with ethanol (vehicle control) or 100 nM E2 for exactly 55 minutes. Media was removed and cells were incubated in CSK buffer at room temperature for exactly 10 minutes. Composition of CSK buffer was as follows: 10 mM PIPES/KOH pH 6.8, 100 mM NaCl, 300 mM sucrose, 1 mM EGTA, 1 mM magnesium chloride, 1 mM DTT, 0.5% Triton-X-100, 1X protease inhibitor cocktail; DTT, Triton-X-100 and protease inhibitor cocktail were added immediately before use. CSK buffer was removed and cells were fixed in 4% paraformaldehyde for 15 minutes at room temperature. After three washes with 1X PBS, cells were incubated in 0.2% Triton solution made in 1% BSA in 1X PBS for 10 minutes at room temperature. Cells were washed thrice with 1X PBS. Coverslips were overlaid with primary antibody mix and incubated in a humidified chamber at 4 degrees Celsius overnight. Antibodies were used at 1:500 dilution in 1% BSA made in 1X PBS. Following antibodies were used: Estrogen Receptor alpha mouse monoclonal antibody (F-10) sc-8002 (Santa Cruz Biotechnology) and anti-Myosin Iβ (Nuclear) rabbit antibody M3567-200UL (Sigma). Coverslips were transferred to 6-well plates and washed thrice with 1X PBST (0.1% Tween-20 in 1X PBS). Fluorophore-conjugated secondary antibodies (A32723 goat anti-mouse Alexa Fluor plus 488, Invitrogen and A32733 goat anti-rabbit Alexa Fluor plus 647, Invitrogen) were used at 1:500 dilution in 1X PBST. Coverslips were overlaid with secondary antibody mix and incubated in a humidified chamber at room temperature in dark. Coverslips were transferred to 6-well plates and washed thrice with 1X PBST. They were incubated for 10 minutes in DAPI solution (0.2 µl in 10 ml PBST) in dark and washed thrice with 1X PBST. Coverslips were mounted on glass slides using 90% autoclaved glycerol and sealed with nail polish. Imaging was performed with Olympus FV3000 six laser confocal microscope. Three biological replicates were imaged.

Image analysis was done with Matlab using custom scripts. Nuclei segmentation was performed using Otsu thresholding followed by filling holes in the binarized masks and removing image border masks. For puncta/foci calling, Unsharp masking was done using Median blur followed by Maxentropy thresholding and watershed segmentation. Finally, foci having a size of 6 or more pixels were selected. Area fraction was calculated as the ratio of area occupied by the foci to the area of the entire nucleus (from nuclear segmentation). Pearson correlation coefficient between ERα and NM1 channels for each nucleus was calculated using the nuclear masks. Significance of difference between medians was tested using two-tailed Mann-Whitney test.

### qPCRs

For RT-qPCRs, cells were harvested in Trizol (15596026; Invitrogen) and stored at - 80 degrees overnight. Samples were thawed to reach room temperature and given a mini spin. Chloroform was added at 1/5^th^ volume, tubes were invert-mixed and left undisturbed for 5 minutes at room temperature. Aqueous phase was collected after centrifugation at 12000 rpm for 12 minutes at 4 degrees Celsius. Equal volume of isopropanol was added and tubes were vortexed and left undisturbed for 10 minutes at room temperature. RNA pellets were obtained by centrifugation at 12000 rpm for 12 minutes at 4 degrees Celsius. The pellets were washed twice with 75% ethanol (made in nuclease-free water) at 12000 rpm for 12 minutes at room temperature. Pellets were air-dried for 15 minutes and dissolved in nuclease-free water. RNA concentration and absorbance ratios were measured with Nanodrop. 1µg RNA was treated with eZDNase (11766051, Invitrogen) and used to prepare cDNA with the help of random hexamers, dNTPs, buffer and DTT from SuperScript IV First-Strand Synthesis System (18091050; Invitrogen) in nuclease-free water as per manufacturer’s guidelines. RNA was kept on ice throughout the protocol and later stored at −80 degrees Celsius. The cDNA thus prepared was diluted 1:10 in nuclease-free water and stored at −20 degrees Celsius. Real time PCR was performed in triplicates with the cDNA and PowerUp SYBR Green Master Mix (A25742, Applied Biosystems) using CFX96 touch real time PCR (Bio-Rad). Primers used for the same are mentioned in table S1. Fold changes in mRNA levels were calculated as 2^-ΔΔCt with GAPDH Ct values for normalization. Paired t-test was performed on data from three biological replicates to test significance.

For ChIP qPCRs, 1 µl of the purified DNA (from input, beads and IP samples) was diluted 10 times in nuclease-free water and used as template. Reactions were set up in triplicates with PowerUp SYBR Green Master Mix (A25742, Applied Biosystems) as per the manufacturer’s guidelines. The reactions were carried out with CFX96 touch real time PCR (Bio-Rad). Primers used for the same are mentioned in table S2. Changes in ERα occupancy were calculated using percent input method. Only those samples that showed enrichment in IP but not beads were used further for ChIP-seq.

### RNA-sequencing and analysis

RNA was isolated as mentioned previously. rRNA depletion was performed using NEBNext® rRNA Depletion Kit V2 (Human/Mouse/Rat) with RNA Sample Purification Beads kit (E7405L). RNA library was made with NEBNext® Ultra™ II Directional RNA Library Prep with Sample Purification Beads (E7765L). Paired-end sequencing was performed on NovaSeq 6000 platform using SP flow cell with 2×50bp sequencing read length. Reads were uploaded to Galaxy web platform and their quality was affirmed using FastQC. Adapters were trimmed with *cutadapt*. Then, the reads were aligned to hg19 with NCBI hg19 RefSeq gtf file as the reference for transcript annotation. Alignment was done with RNA STAR on Galaxy with following parameters: transcript (--sjdbGTFfeatureExon); 50 (--sjdbOverhang); TranscriptomeSAM GeneCounts(-- quantMode); WithinBAM –chimOutType; remaining parameters were kept at default. Using *MarkDuplicates* and *Filter BAM*, duplicates were marked and removed, and the output BAM files were filtered. Only reads that were mapped with high mapping quality (>=20 phred score), were properly paired and were primary alignments were retained. BAM files were sorted with *Samtools sort* and used as input for *htseq-count.* Counts per gene were obtained using the following parameters in *htseq-count*: Union (-- mode); Reverse (--stranded) [as the libraries were reverse stranded]; Transcript (-- type); All (--nonunique). hg19 NCBI RefSeq gtf file was used as the reference. Secondary alignments were not included and other parameters were left at default. The resultant count files from both biological replicates were analyzed for differential gene expression using DESeq2. Induction of E2 response was ascertained by comparing vehicle-treated and E2-treated samples. siScr and siMyo1c samples (both +E2) were compared to check the effect of Myo1c knockdown on gene expression. The corresponding result files from DESeq2 were used to construct volcano plots in Galaxy server with significance threshold set at 0.05.

Further, normalized gene expression values were calculated as transcripts per million (tpm). Briefly, the counts per gene in the *htseq-count* files were normalized to gene length to obtain reads per kilobase (rpk). Then a scaling factor to normalize for sequencing depth was calculated using all the rpk values in a sample. The rpk was divided by the scaling factor to obtain tpm for each gene. Next, genes classified as upregulated from GRO-seq after 40 minutes of E2 treatment in MCF-7 cells by Hah et al. (2011) were chosen and their log2 tpm was calculated in siScr and siMyo1c samples (+E2). Low-expressing genes (log2 tpm < 0.5) were filtered out. Significance of difference between medians was tested using two-tailed Mann-Whitney test.

For obtaining counts of intron-mapped transcripts, NCBI hg19 RefSeq gtf file was converted to BED12 format and intron list was extracted with *Gene BED To Exon/Intron/Codon BED expander* tool in Galaxy server. The intron list was sorted by chromosome and start position with *SortBed*. The identifier ‘transcript’ was added in the third column, start and end position values were moved to the fourth and fifth columns respectively and individual accession numbers were removed, while retaining rest of the information in the BED file. Then, using *bedtools MapBed*, the information in a sorted hg19 NCBI RefSeq gtf file was mapped onto the intron list BED file. The resultant gtf file was used in *htseq-count* along with the filtered, sorted BAM files obtained from RNA-seq to get counts of intron-mapped transcripts per gene. *htseq-count* parameters were the same as mentioned above.

### Data analysis and visualization

Western blot images were visualized and quantified using Fiji (Schindelin et al., 2012). ChIP-seq heatmaps were generated with Galaxy server (Jalili et al., 2020). Integrated Genomics Viewer (Robinson et al., 2012) was used to visualize ChIP-seq tracks. Confocal microscopy images were visualized with Fiji. GraphPad Prism 8.0.2 for Windows was used to perform tests of significance and plot the data. Schematics were designed in BioRender.

## Acknowledgements

We acknowledge the support of the NCBS-TIFR under the Department of Atomic Energy (Government of India) no. RTI 4006 (to DN). DN is an EMBO Global Investigator (GIN). We also acknowledge the funding support from Welcome-IA (IA/S/23/1/506749). SS and AHN are thankful to the Council of Scientific & Industrial Research (CSIR, India) for providing Shyama Prasad Mukherjee Fellowship and graduate research fellowship respectively. RM and DS acknowledge graduate research fellowship provided by NCBS-TIFR. We are grateful to Awadhesh Pandit and Lakshminarayanan CP affiliated with the Next Generation Sequencing Facility at NCBS for help with NGS experiments. Confocal microscopy was done at Central Imaging and Flow Cytometry facility at NCBS. We thank Dr. Minhaj Sirajuddin and Nivya Mendon for discussion. We thank all members of DN lab for helpful suggestions and comments on the manuscript.

## Author contribution

SS, IAP and DN designed the study. SS and IAP performed the experiments with help from DS. SS analyzed the biochemical and genomics data. AHN performed confocal imaging. RM analyzed the microscopy data. SS and DN wrote the manuscript. All authors participated in editing the manuscript.

---

**Figure S1.**
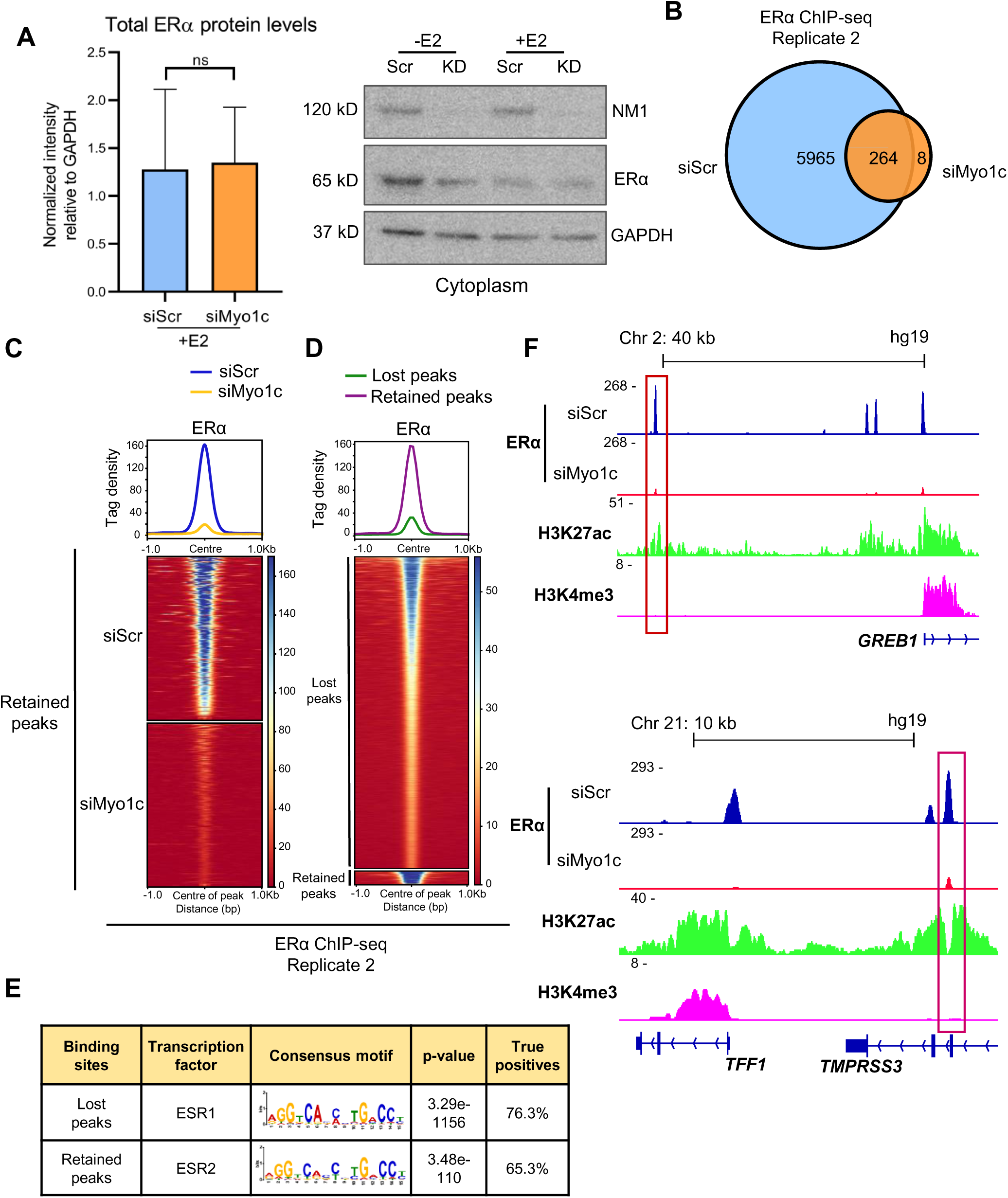
Genomic occupancy of ERα is dependent on Myo1c. (A) Bar graph showing total cellular ERα protein levels do not change upon Myo1c knockdown (left panel). Background-normalized band intensities of ERα relative to those of GAPDH are plotted. Paired t-test was used to test significance (p-value=0.908); whiskers indicate standard deviation around mean; N=3. Western blot showing ERα translocation into nuclei upon E2 stimulation is unaffected by Myo1c knockdown; -E2: vehicle control; +E2: E2 treatment (middle panel). (B) Venn diagram showing ERα peaks detected in second biological replicate of ERα ChIP-seq in siScr and siMyo1c samples at 60 minutes of E2 stimulation. (C) Heatmap showing ERα ChIP-seq signal in the second replicate decreases even at the retained peaks in the siMyo1c sample, as compared to siScr sample. (D) Heatmap showing ‘retained’ peaks have higher mean ERα signal compared to ‘lost’ peaks. ERα signal intensity was plotted using second biological replicate of siScr sample. (E) Table showing motifs with highest enrichment score in the different categories of peaks shown in panel B. (F) IGV screenshots showing loss of ERα ChIP-seq signal upon Myo1c knockdown (second biological replicate) at enhancer clusters regulating *GREB1* gene and *TFF1* gene. Major enhancers are highlighted in red boxes. All tracks show ChIP signal from MCF-7 cells post 60 minutes of E2 stimulation; H3K27ac and H3K4me3 tracks were generated using published datasets (see Methods).

**Figure S2:**
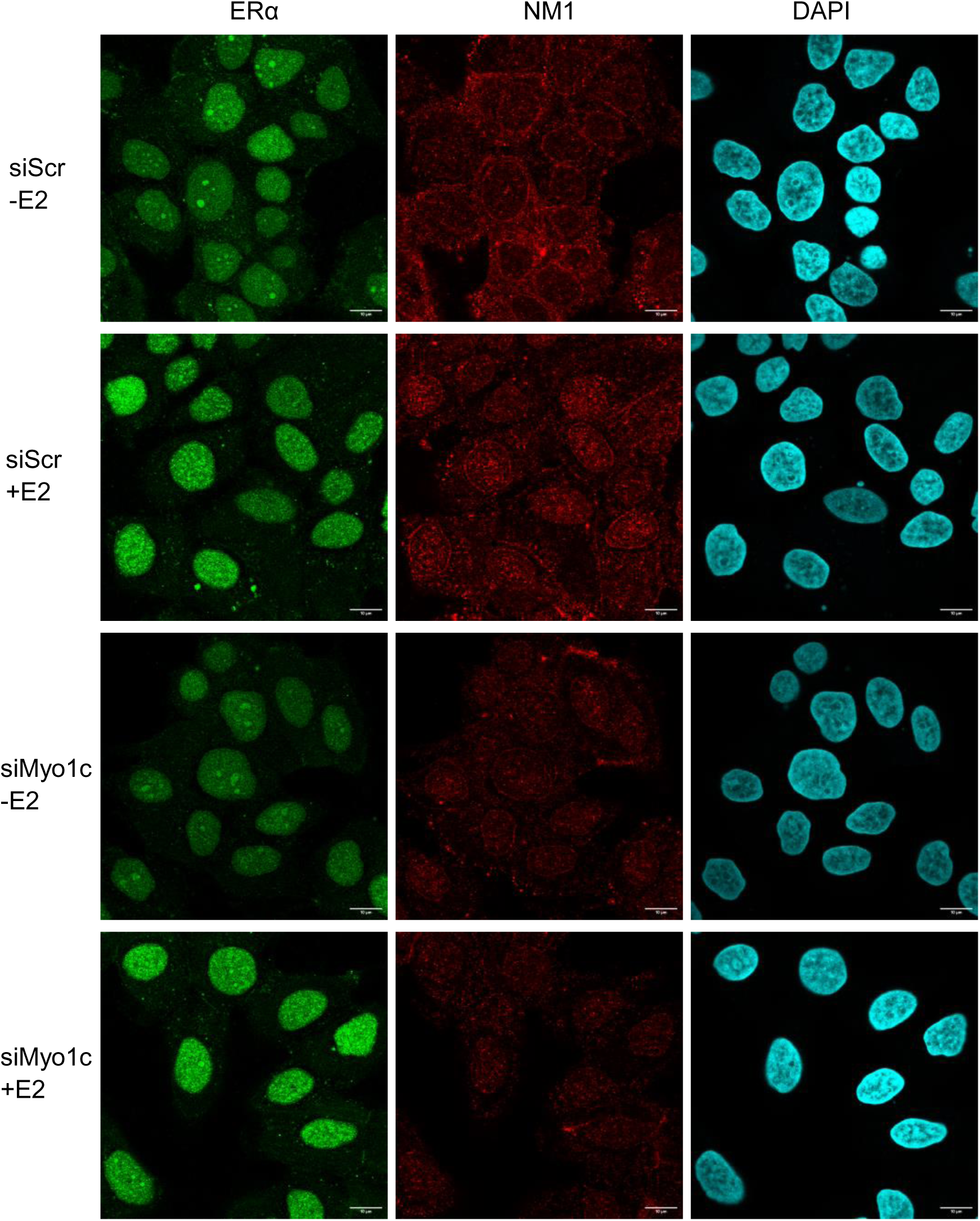
Myo1c depletion perturbs ligand-dependent ERα condensates. Confocal images of immunofluorescence for ERα (green) and NM1 (red) in CSK-treated samples. Both NM1 and ERα show an increase in nuclear localization post 60 minutes of E2 treatment. Knockdown of Myo1c is evident from the overall reduction in NM1 signal in the siMyo1c samples. DAPI is shown in cyan. Scale bar: 10 μm. -E2: vehicle treatment for 60 minutes; +E2: E2 treatment for 60 minutes.

**Figure S3:**
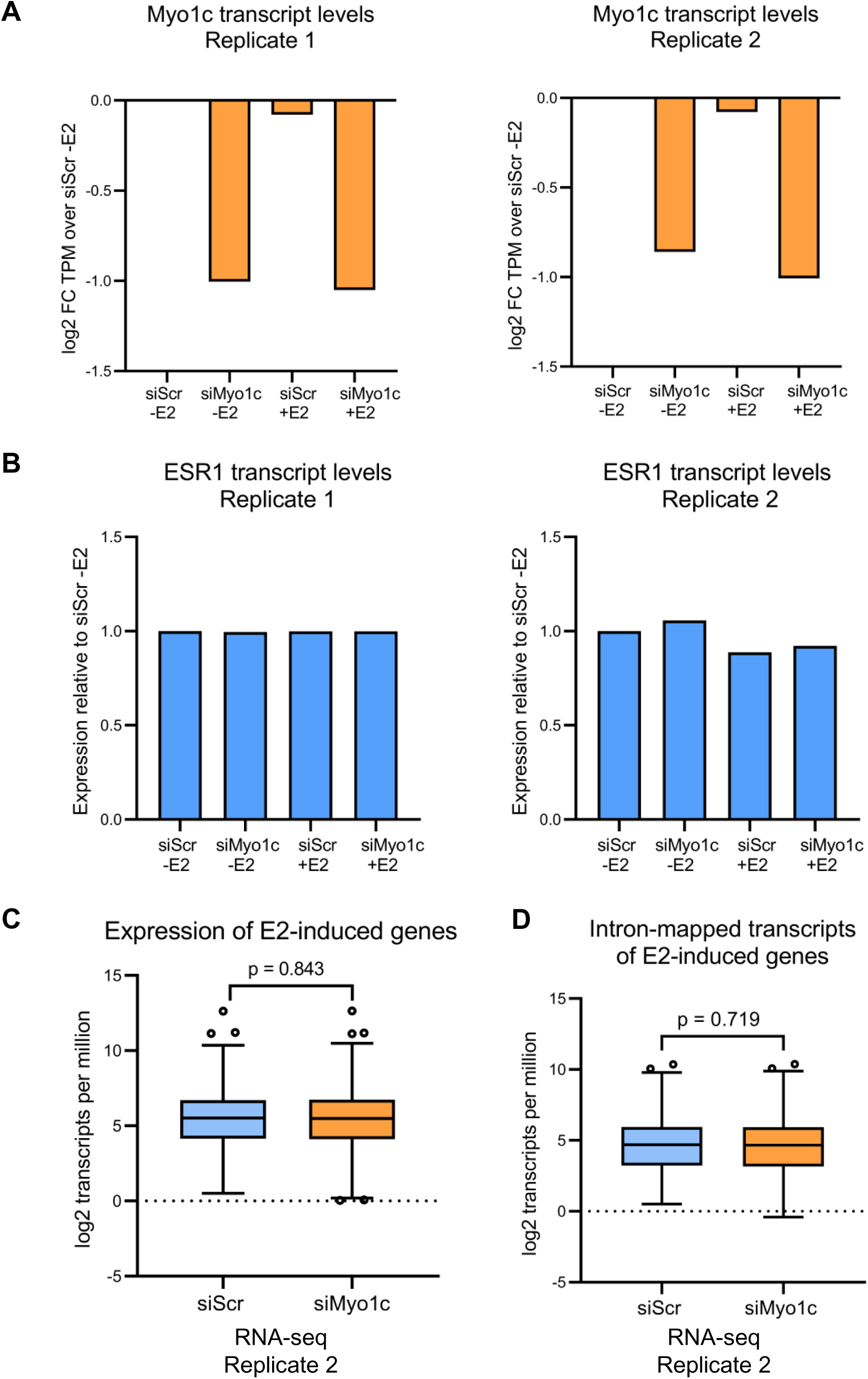
Transcription of E2-induced genes is unaffected by Myo1c knockdown. (A) Confirmation of Myo1c knockdown in different samples. Values plotted are log2 fold change of *MYO1C* transcripts per million in different samples relative to siScr sample treated with vehicle control for 60 minutes (siScr -E2). (B) *ESR1* gene transcription is not affected by Myo1c knockdown. Values plotted are ratio of *ESR1* transcripts per million in each sample to that in siScr -E2 sample. (C) Steady-state transcript levels of 1200 top E2-induced genes (Hah et al., 2011) from second biological replicate of total RNA-seq. (D) Quantification of transcripts of E2-induced genes (same as panel C) mapping to introns from second biological replicate of total RNA-seq. P-values in (C) and (D) were calculated using Mann-Whitney test. Centre lines within boxes represent medians. Whiskers represent values within 1.5 IQR (Tukey).

**Table S1.**
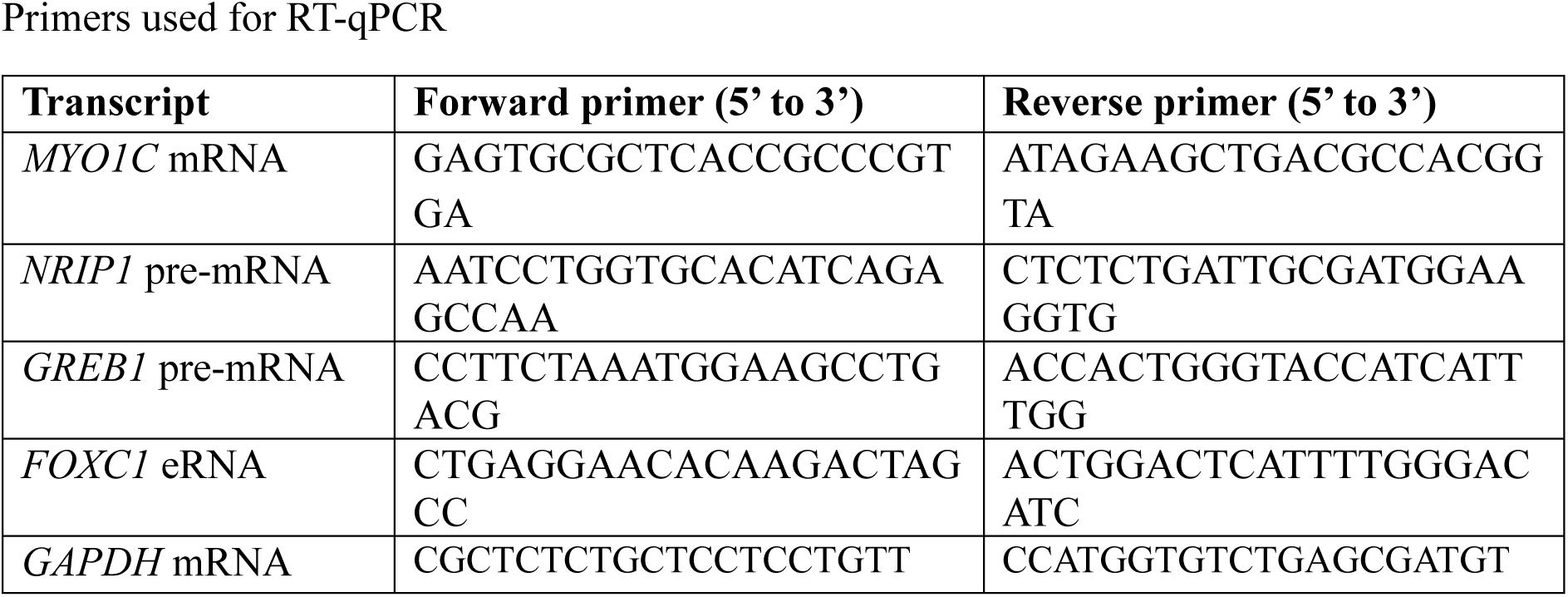
Primers used for RT-qPCR.

**Table S2.**
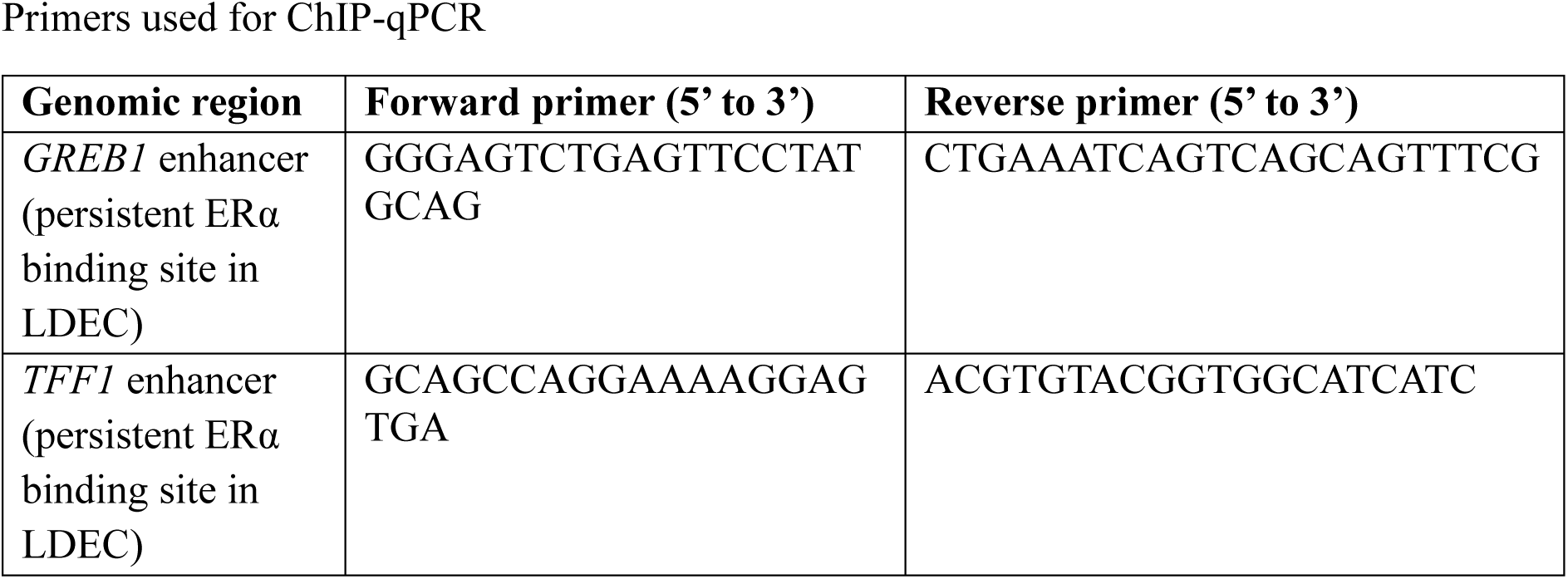
Primers used for ChIP-qPCR.

## Notes

### Competing Interest Statement

The authors have declared no competing interest.

